# Discovery of two potential new species and two novel bat-coronavirus subgenera (*Phyllacovirus* and *Phyllobecovirus*) in the Neotropics

**DOI:** 10.64898/2026.09.23.753618

**Authors:** Leonardo Corrêa da Silva Junior, Filipe Zimmer Dezordi, Alexandre Freitas da Silva, Jennifer Oliveira Melo, Yago José Mariz Dias, Paola Cristina Resende, Luciana Reis Appolinario, Marilda Siqueira, Maria Ogrzewalska, Gabriel da Luz Wallau

**Author notes:** Both authors contributed equally to this work (Leonardo Corrêa da Silva Junior, Filipe Zimmer Dezordi).

## Abstract

Bats are major natural reservoirs for coronaviruses, yet complete viral genomes from South America remain scarce, limiting evolutionary and taxonomic understanding. Here, we conducted metatranscriptomic sequencing of coronavirus-positive bat samples collected across two ecologically distinct Brazilian biomes: the Atlantic Forest and the semi-arid Caatinga. We recovered seven complete or near-complete genomes belonging to *Alphacoronavirus* and *Betacoronavirus*. Phylogenetic and comparative similarity analyses of conserved replicase domains (3CLpro, NiRAN, RdRp, ZBD, HEL1), following International Committee on Taxonomy of Viruses (ICTV) demarcation criteria, revealed significant viral diversity. Within *Alphacoronavirus*, two genomes from Atlantic Forest phyllostomid bats (*Artibeus lituratus* and *Carollia perspicillata*) formed a deeply divergent sister lineage to *Amalacovirus*, exhibiting a mean amino acid similarity of 76.7% with the reference genome. Within *Betacoronavirus*, one genome from a Caatinga phyllostomid bat (*Artibeus planirostris*) clustered within the recently described *Ambecovirus* clade, displaying 75.9% mean amino acid similarity with mormoopid-associated reference sequences. Based on these divergence levels and non-recombinant genomic architectures, we propose two novel candidate subgenera, *Phyllacovirus* and *Phyllobecovirus*, alongside potential novel viral species. Furthermore, our findings demonstrate strong host-associated structuring and biogeographical partitioning of viral lineages across Neotropical biomes. Overall, this study expands the genomic landscape of South American bat coronaviruses and underscores the importance of continuous genomic surveillance at human-wildlife interfaces.

## 1. Introduction

Coronaviruses (CoVs) comprise a diverse group of positive-sense single-stranded RNA viruses belonging to the family *Coronaviridae*, order Nidovirales (Anthony et al., 2017). They are classified into four genera, *Alphacoronavirus* (AlphaCoV), *Betacoronavirus* (BetaCoV), *Deltacoronavirus* (DeltaCoV), and *Gammacoronavirus* (GammaCoV), and are broadly distributed among birds and mammals (Woo et al., 2012). While DeltaCov and GammaCoV primarily infect avian hosts, they can also infect certain mammals; notably, porcine DeltaCoV has recently been identified as a causative agent of human infections (Lednicky et al., 2021). In contrast, AlphaCoV and BetaCoV predominantly infect mammals (Zhu et al., 2020). Although these genera can infect a wide range of hosts, bats harbor an exceptional diversity of CoVs and are considered the natural reservoirs for the majority of lineages (Woo et al., 2006, 2012).

CoVs are also known to spill over between species, crossing the species barrier and eventually adapting to new hosts, which can lead to outbreaks (Caraballo, 2022). More prominent examples are the three Covs that caused transitory and pandemic outbreaks in humans such as SARS-CoV, MERS-CoV and SARS-CoV-2 (Zhou et al., 2020), but spill over to other mammals leading to outbreaks are also known including an alphacoronavirus from bats to pigs (Zhou et al., 2018) and a deltacoronavirus from birds to pigs (Kong et al., 2022). Reflecting this evolutionary dynamic, bats harbor the largest diversity of CoVs among mammals, with AlphaCoV being more widespread and abundant than BetaCoV (Wong et al., 2019). For example, recent analyses have identified several subgenera within AlphaCoVs (such as *Nyctacovirus*, *Myotacovirus*, *Decacovirus*, and *Minunacovirus*), as well as lineages that remain undescribed, whereas the genus BetaCoVs comprises a more limited number of well-defined subgenera (Rieu et al., 2026). Nevertheless, a new BetaCoVs subgenus named *Ambecovirus* has been recently identified in South American bats suggesting that our understanding of the full diversity of this genus is still expanding (Wallau et al., 2025).

Furthermore, ecological surveys indicate a high frequency of *Alphacoronavirus* detection in bat populations. These findings suggest that certain *Alphacoronavirus* may exhibit widespread circulation and a high capacity for transmission among individuals within the same population (Mombo et al., 2026). An increasing number of studies have demonstrated that bats serve as natural and ancestral reservoirs for both AlphaCoVs and BetaCoVs, playing a central role in maintaining viral lineages that are phylogenetically related to those that have sporadically emerged in humans (Bueno et al., 2022). The detection of multiple coronavirus lineages in bats occurs across a wide range of host species, reflecting the broad ecological, taxonomic, and behavioral diversity of bats, which occupy numerous ecological niches and play essential roles in ecosystem functioning (Luis et al., 2013; Jones et al., 2023). Expanding research focused on bat-associated coronaviruses is critical for understanding evolutionary, ecological, and epidemiological processes underlying the maintenance, emergence and reemergence of these important viral group carrying several zoonotic pathogens (Weber and Da Silva, 2023).

Bat-associated coronaviruses in the Americas exhibit high genetic diversity and a wide geographic distribution, reflecting a long evolutionary history between these viruses and their hosts (Ruiz-Aravena et al., 2022). Brazil hosts one of the richest bat faunas in the world, with 186 species recorded across nine families distributed throughout multiple biomes (Garbino et al., 2024). Despite this remarkable diversity, viral surveillance studies in Brazilian bats remain limited due to no systematic longitudinal sampling, limited host coverage and unequal sampling through different biomes (Letko et al., 2020; Wallau et al., 2023). Most studies remain restricted to short RdRp fragments, and an exploratory analysis of the NCBI Virus database suggests that only approximately 3.9% of publicly available bat coronavirus sequences correspond to complete genomes (NCBI, 2026). Among the available records, partial sequences of approximately 500 bp from Alpha- and Beta-CoVs predominate, obtained from bats inhabiting different regions and belonging to the families Molossidae, Vespertilionidae, and Phyllostomidae (Bittar et al., 2020; Lima et al., 2013; Góes et al., 2013, 2016). Furthermore, *Desmodus rotundus* from the Pampa biome has been reported to harbor AlphaCoVs (Alves et al., 2022), while another positive sample was identified in bats from the Atlantic Forest biome. BetaCoVs have also been detected in bats of the family Mormoopidae in the Atlantic Forest biome, in southeastern Brazil (Brandão et al., 2008).

Although bats harbor a remarkable diversity of CoVs, complete genomes remain scarce in public databases and the scientific literature. This genomic gap is particularly evident in South America, where complete bat coronavirus genomes have been reported in only a few geographic and ecological contexts, including an AlphaCov from *Tadarida brasiliensis* in the Argentine Pampas (Cerri et al., 2023), an AlphaCoV from *D. rotundus* in the Peruvian Amazon biome (Bergner et al., 2020), and two BetaCoVs genomes from *Pteronotus gymnonotus* associated with the Caatinga and Atlantic Forest biomes in northeastern Brazil (Wallau et al., 2025). Taken together, these findings highlight the significant scarcity of available complete genomic data on bat-associated coronaviruses in South American ecosystems. Obtaining complete genomes is essential for robust phylogenetic analyses, for identifying recombination events, and for clarifying evolutionary relationships among strains originating from different hosts and geographic regions (Bergner et al., 2020).

Despite the current scarcity of complete genomes and the historical reliance on these short RdRp sequence fragments, some interesting findings have been drawn from the available data.These show that alphacoronaviruses are more predominantly detected in bats from Brazil and the wider Americas region than betacoronaviruses, likely reflecting the higher diversity (more subgenera) and worldwide prevalence of the former (Ruiz-Aravena et al., 2022). And that alphacoronaviruses are more prone to cross-species transmission (CST), evolving through a combination of cospeciation and CST, than betaconronaviruses that cospeciate more frequently showing little evidence of CSTs so far (Caraballo, 2022; Wallau et al., 2025). Additionally, considering the main bat host families, CSTs of alphacoronaviruses of at least four different lineages were detected between Phyllostomidae species, while a lower number of events were also found between species of Vespertilionidae and Molossidae families underlying the important role of Phyllostomidae species in short and long-term evolution of Alphacoronavirus in the Americas (Caraballo, 2022). More recent genomic findings also uncovered recombination events in the spike protein of alphacoronaviruses found in *T. brasiliensis* highlighting the still little explored evolutionary phenomena shaping the diversity and adaptation of AlphaCoVs in the Americas (Cerri et al., 2023).

In this study, we performed the complete genomic characterization, through metatranscriptomic sequencing, of bat samples collected from two ecologically distinct Brazilian biomes, the Atlantic Forest and the Caatinga, which had previously been tested and found positive for coronaviruses by Sanger sequencing. Samples from the Atlantic Forest were previously identified as positive for *Alphacoronavirus* by da Da Silva Junior et al. (2025), whereas samples from the Caatinga were identified as positive for *Alphacoronavirus* and *Betacoronavirus* by Figueiroa et al. (2025). Thus, whole-genome sequencing was employed as a complementary approach to the previously performed molecular characterization, with the aim of increasing the genomic resolution of these viruses. The generation of draft and complete genomes enabled a more comprehensive assessment of genetic diversity, including the identification and analysis of mutations across the genome, the investigation of potential recombination events, and a more robust phylogenetic characterization of the identified lineages. Of note, the genomic data generated supported a proposal of a novel CoV species and two new subgenera expanding our understanding of CoVs diversity and evolution in the neotropics.

## 2. Results

**Table 1.** Genomes generated in this study. ^1^Number of non-sequenced bases when compared with the closest reference genome used for scaffolding; ^2^Million reads mapped. ^3^Mean coverage depth; ^4^Host species. ^5^Collection date. All samples are collected by rectal swab.

| Sequence | Clade (subgenus) | Length | N<sup>1</sup> | M reads<sup>2</sup> | Depth<sup>3</sup> | rpkm | Species<sup>4</sup> | Date<sup>5</sup> | State |
| --- | --- | --- | --- | --- | --- | --- | --- | --- | --- |
| FMA520 | *Phyllacovirus* | 28385 | 10983 | 0.105 | 188.57 | 35229.87 | *Artibeus lituratus* | 2021-02-09 | RJ |
| FMA536 | *Phyllacovirus* | 30681 | 204 | 23.35 | 38482.47 | 32593.46 | *Carollia perspicillata* | 2021-03-15 | RJ |
| NAC28 | *Amalacovirus*-like | 29097 | 3953 | 0.113 | 194.26 | 34367.80 | *Carollia perspicillata* | 2022-03-22 | CE |
| NAC60 | *Amalacovirus* | 32258 | 4931 | 0.082 | 130.64 | 31000.06 | *Lonchorhina aurita* | 2022-03-19 | CE |
| PC112 | *Amalacovirus* | 29540 | 5085 | 0.051 | 87.51 | 33852.40 | *Glossophaga soricina* | 2022-03-13 | CE |
| PC122 | *Amalacovirus* | 29346 | 1270 | 0.148 | 255.43 | 34076.19 | *Phyllostomus discolor* | 2022-03-15 | CE |
| NAC39 | *Phyllobecovirus* | 29539 | 0 | 77.07 | 132357.5 | 33853.55 | *Artibeus planirostris* | 2022-03-18 | CE |

Sequences derived from six distinct samples clustered with *Alphacoronavirus* from the *Amalacovirus* subgenus, with high branch support (aLRT = 91.6; ultrafast bootstrap = 99), and were associated with viruses previously identified in bats of the family Phyllostomidae, the same host family from which the samples were collected (Figure 1A).

**Figure 1:**
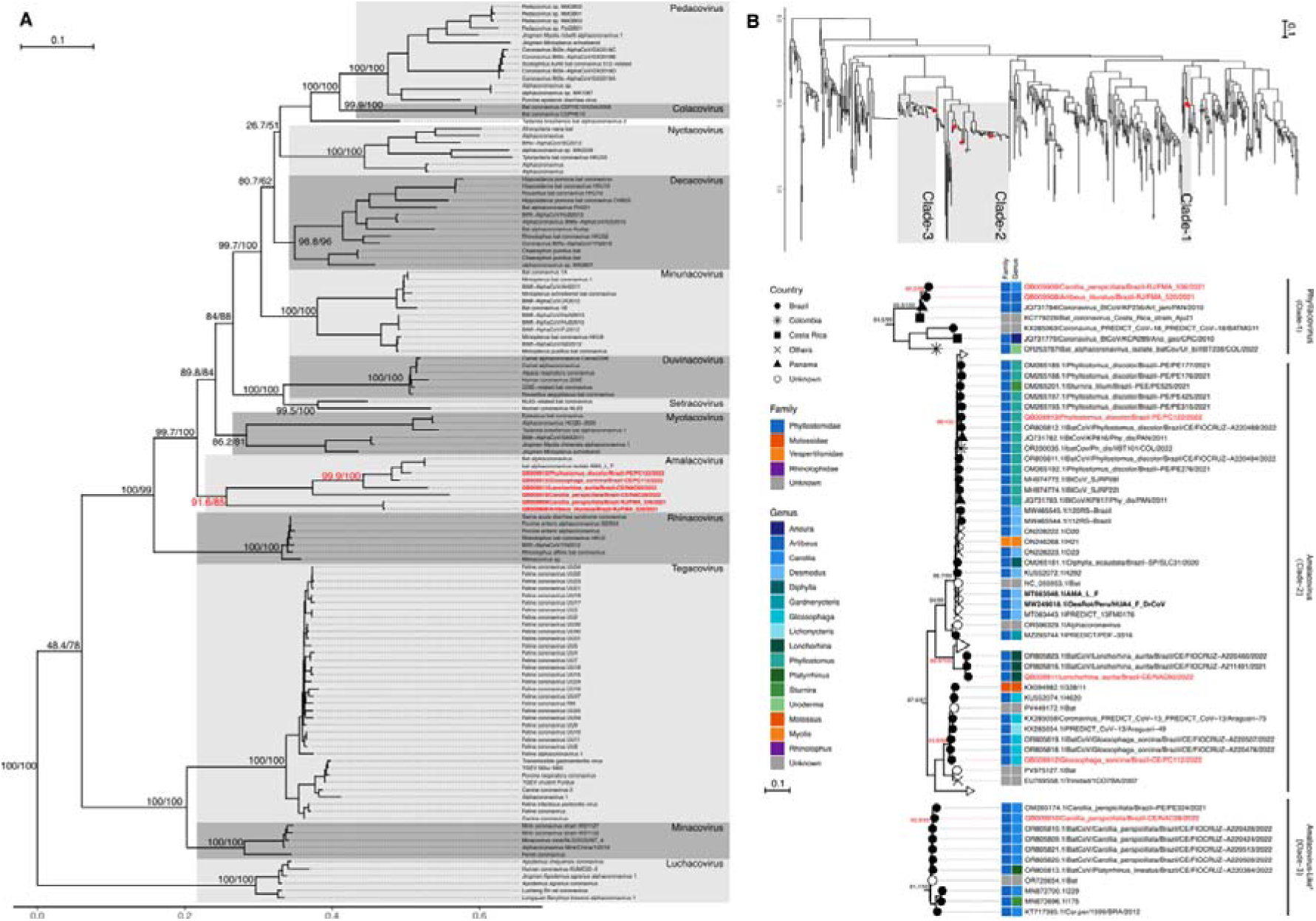
*Alphacoronavirus* phylogeny. **A.** Maximum likelihood phylogenetic reconstruction of *Alphacoronavirus* genus based on amino acid sequences of conserved domains (3CLpro, NiRAN, RdRP, ZBD, and HEL1 domains, 1960 residues). **B.** Maximum likelihood phylogenetic reconstruction of *Amalacovirus* subgenus based on the RdRp coding region including partial (446 residues). Genome sequenced in this study are marked in red. Branch support informed as aLRT/Ultrafastboostrap. Bold black sequences represent the complete genomes of *Amalacovirus* described in the literature. * A sister clade of *Amalacovirus* (Clade-3) described in different studies (Góes et al., 2016; Anthony et al., 2017; Bueno et al., 2022). Sequence names with accession number and sample code only had no extra information available.

Phylogenetic analysis of 446-nucleotide RdRp fragments from *Amalacovirus* and *Amalacovirus-*like viruses recovered multiple clades (Figure 1B). Genomes from samples FMA520 and FMA536 clustered outside the *Amalacovirus* and *Amalacovirus*-like clades, grouping instead with other unclassified *Alphacoronavirus* sequences, with high node support (99.2/98 for aLRT/ultrafast bootstrap - Figure 1B - clade I). The genome from sample NAC28 clustered within the *Amalacovirus*-like clade (92.9/99 support) alongside other Brazilian viruses isolated from the same host species (*Carollia perspicillata*). In contrast, genomes from samples NAC60, PC112, and PC122 clustered within the *Amalacovirus* clade, forming distinct subclades that reflected host-associated divergence. Despite differences in sequence length and analytical strategy, both phylogenetic reconstructions consistently indicate that genomes from samples FMA520 and FMA536 are closely related to *Amalacovirus* but may represent a novel sister subgenus. No recombination signal was identified on FMA520 and FMA536 genomes.

Phylogenetic analysis of the genome from sample NAC39 revealed the presence of a previously unreported *Ambecovirus* (Figure 2A), which clustered with viruses detected from the same host genus (*A. planirostris* and *A. lituratus*, Figure 2B). For the alphacoronavirus assigned near to the subgenus *Amalacovirus*, the of FMA_536 shared 60.73% nucleotide similarity with Bat alphacoronavirus isolate AMA_L_F (MT663548.1, Figure 3A). At the amino acid level, over 1,947 residues of RdRp conserved domains of FMA_536 and FMA_520 were nearly identical to one another (99.02%), while sharing 77.08% and 76.37% similarity with the AMA_L_F reference (QLE11824.1, Figure3B), respectively (mean 76.7%). For the betacoronavirus assigned to the subgenus *Ambecovirus*, NAC39 shared a mean of 62% nucleotide similarity with two *Pteronotus gymnonotus* genomes from Pernambuco (N107_15 and N107_12, Figure 3C). NAC39 showed 79% and 80% amino acid similarity (mean 79.5%) with the corresponding *P. gymnonotus* sequences (N107_10 and N107_12) over 1,893 residues of RdRp conserved domains (Figure 3D).

**Figure 2:**
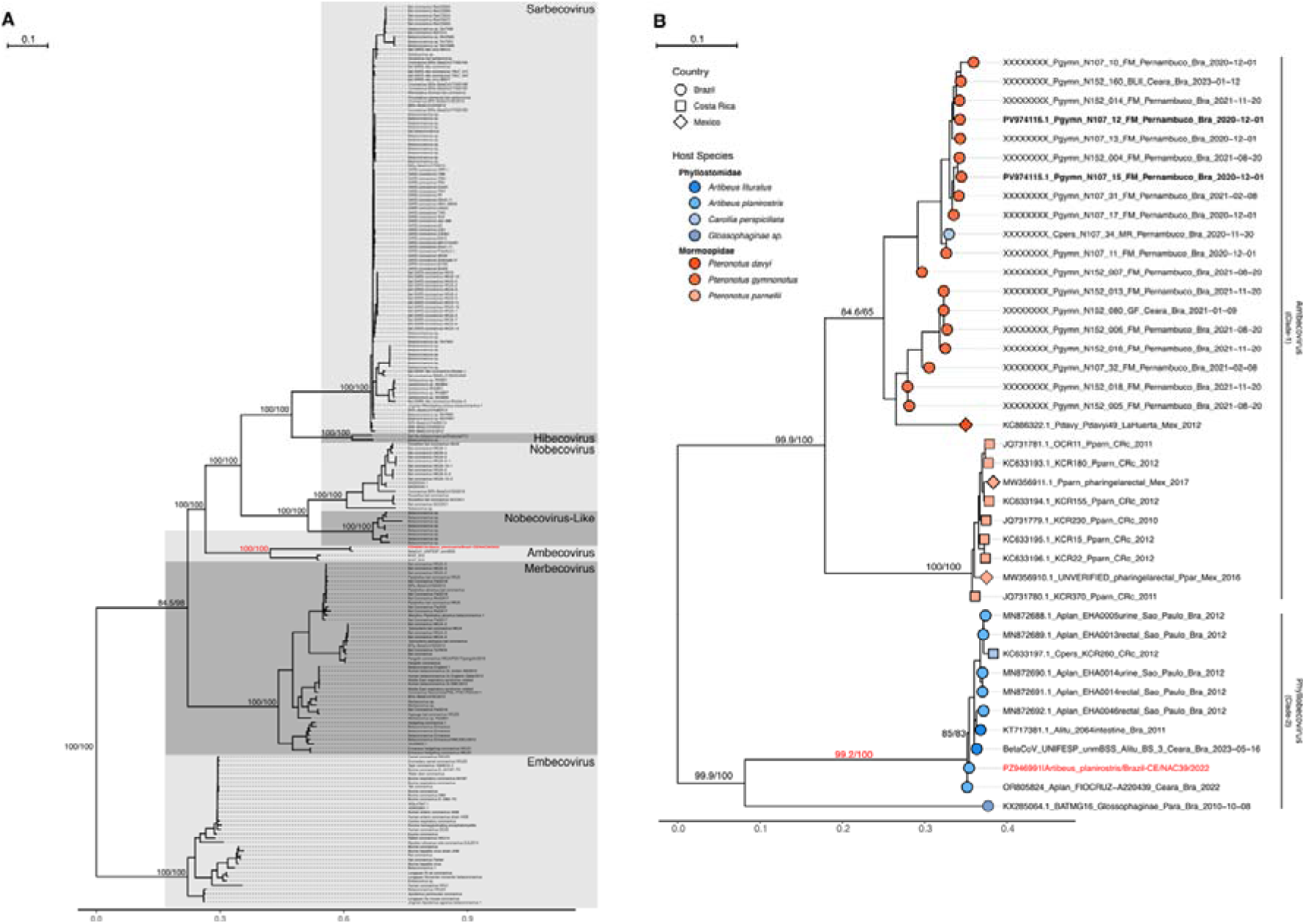
*Betacoronavirus* phylogeny. **A** - Maximum likelihood phylogenetic reconstruction of Betacoronavirus genus based on amino acid sequences of conserved domains (3CLpro, NiRAN, RdRP, ZBD, and HEL1 domains, 1978 residues). **B** - Maximum likelihood phylogenetic reconstruction of *Ambecovirus* subgenus based on the RdRp coding region including partial (439 bp) and full RdRp nucleotide coding region (2790 residues) based on *Ambecovirus* draft genomes informed on Wallau *et al*. (2025). Bold black sequences represent the complete genomes of *Ambecovirus* described in the literature. Genome sequenced in this study are marked in red. Branch support informed as aLRT/Ultrafastboostrap.

**Figure 3:**
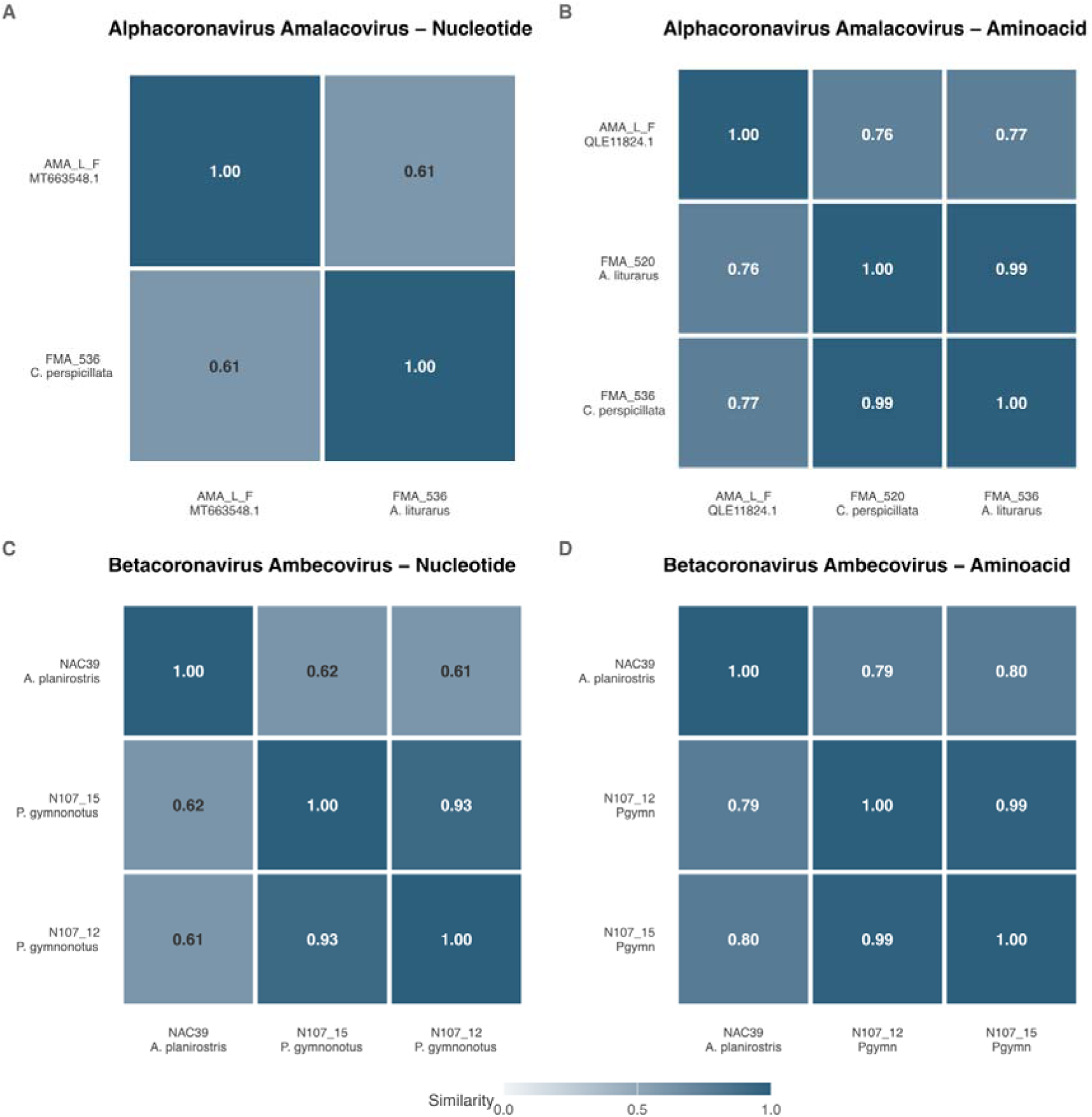
Distance matrices. **A** - Considering the entire genome of *Amalacovirus*. **B** - Considering 3CLpro, NiRAN, RdRP, ZBD, and HEL1 domains of *Amalacovirus*. **C** - Considering the entire genome of *Ambecovirus*. **D**. Considering 3CLpro, NiRAN, RdRP, ZBD, and HEL1 domains of *Ambecovirus*.

## 3. Discussion

The recovery of seven bat coronavirus genomes sampled across two ecologically distinct and biogeographically key Brazilian biomes, the highly biodiverse yet threatened Atlantic Forest, and the semi-arid Caatinga, revealed substantially greater viral diversity than previously inferred from studies based on short fragments of the RdRp gene (NCBI, 2026). Although previous surveys have consistently reported a predominance of alphacoronavirus among Neotropical bats (Caraballo, 2022; Ruiz-Aravena et al., 2022), most available sequences lacked sufficient phylogenetic resolution to identify deeply divergent lineages or resolve higher-level evolutionary relationships. In this context, the simultaneous characterization of viruses belonging to the *Amalacovirus*, *Amalacovirus*-like, and *Ambecovirus* clades, together with two previously unclassified alphacoronaviruses, suggests the existence of a cryptic viral diversity that remains largely unexplored in South America.

Phylogenetic reconstructions revealed that the recovered viruses represent distinct levels of evolutionary divergence within both *Alphacoronavirus* and *Betacoronavirus*. Particularly noteworthy was the phylogenetic placement of samples FMA520 and FMA536, which consistently clustered as a sister lineage to the *Amalacovirus* and *Amalacovirus*-like clades without being clearly assigned to either group. The stability of this pattern across different analyses suggests the existence of a deeply divergent evolutionary lineage that has not yet been formally recognized within current coronavirus taxonomy. Here, we formally propose that the genomes FMA520 and FMA536 represent a novel viral species within a newly proposed candidate subgenus in the *Alphacoronavirus* genus, named *Phyllacovirus*. Following the official ICTV nomenclature guidelines, this name is constructed using the host-derived prefix ’Phyll’, highlighting its restricted evolutionary association with bats of the family Phyllostomidae, followed by the genus suffix ’acovirus’. In support of this taxonomic assignment, comparative similarity analyses using the conserved replicase domains (3CLpro, NiRAN, RdRp, ZBD, and HEL1), which represent the official ICTV demarcation standard for the *Orthocoronavirinae* subfamily, revealed that FMA520 and FMA536 share a high amino acid identity of 99.02% (over 1,947 residues) with each other, demonstrating they belong to the same species. However, they share a mean of only 76.7% amino acid similarity (77.08% for FMA536 and 76.37% for FMA520) with the only available complete reference genome for *Amalacovirus* (Bat alphacoronavirus isolate AMA_L_F, MT663548.1). At the nucleotide level, FMA536 shares only 60.73% similarity with the AMA_L_F reference. Genetic divergence values of this magnitude fall well below the established thresholds for sharing a subgenus or species within *Orthocoronavirinae*.

This taxonomic distinction is further supported by the complete absence of recombination signals in FMA520 and FMA536 genomes. Furthermore, the detection of this novel non-recombinant lineage in two distinct host genera of the Phyllostomidae family, *A. lituratus* (FMA520) and *C. perspicillata* (FMA536) in Rio de Janeiro, suggests that this candidate subgenus is ecologically stable and capable of cross-species transmission (CST) within New World phyllostomid bats. Although additional analyses based on ICTV taxonomic criteria will be necessary to clarify its classification status, this finding expands our understanding of *Alphacoronavirus* radiation in Neotropical bats and suggests that the evolutionary diversity of these viruses remains underestimated. This underestimation is further illustrated by the recent characterization of novel divergent *Alphacoronavirus* lineages in Central America, such as the novel taxa PYCoV4 (detected in *Sturnira parvidens*) and PYCoV7 (detected in the insectivorous *Lophostoma brasiliense*) in Mexico (Jiménez-Rico et al., 2026), which mirror the high cryptic diversity and host-associated structuring we observed in Brazilian biomes. Similar patterns have been reported in studies from Asia and Africa, where increased genomic sampling led to the identification of novel lineages and subgenera within *Alphacoronavirus* (Woo et al., 2012; Rieu et al., 2026).

The identification of a new genome belonging to the recently suggested *Ambecovirus* group represents a particularly important contribution to our understanding of *Betacoronavirus* evolution in the Americas, where this group remains poorly represented in public databases. In this context, the recovery of genome NAC39 (from *A. planirostris* in Ceará) expands the currently known diversity of this group and provides an opportunity to investigate evolutionary patterns that were previously inferred from a restricted dataset. Here, we formally propose that NAC39 represents a novel viral species within a candidate subgenus named *Phyllobecovirus*, establishing a symmetrical, host-associated taxonomy alongside *Phyllacovirus* for Neotropical bat coronaviruses. Conventionally following the official ICTV nomenclature guidelines, the name *Phyllobecovirus* is constructed by combining the host family prefix ’Phyllo’ (Phyllostomidae) with the *Betacoronavirus* genus suffix ’becovirus’. This taxonomic designation is strongly supported by comparative genomic analyses of the conserved replicase domains (3CLpro, NiRAN, RdRp, ZBD, and HEL1), which reveal that NAC39 shares a mean of 79.5% amino acid identity (80% with N107_15 and 79% with N107_12 over 1893 residues) and a mean of 62% nucleotide similarity with the *Pteronotus gymnonotus*-associated sequences from Pernambuco (N107_10 and N107_12). A genetic divergence of approximately 20% in these highly conserved replicase domains falls well below the established thresholds for sharing a species or subgenus designation within *Orthocoronavirinae*, thereby justifying the formal demarcation of *Phyllobecovirus* as a distinct, ecologically stable subgenus representing the first complete genomic characterization of a phyllostomid-associated lineage in this group.

Unlike the pattern observed among alphacoronaviruses, which are characterized by multiple host-switching events and frequent shifts among host taxa, the currently available evidence suggests that Ambecovirus exhibits much more restricted host associations. While the Pernambuco N107 lineages are associated with mormoopid bats (*Pteronotus* spp., Clade I), NAC39 clusters strictly with clade II lineages previously detected in phyllostomid hosts (*A. planirostris* and *A. lituratus*). This strict partitioning suggests that host association and ecological conservatism have contributed to the diversification of this subgenus, resulting in phylogenetic patterns that are more closely associated with host evolutionary history (Caraballo, 2022; Wallau et al., 2025). This highly conserved, host-restricted pattern contrasts sharply with the evolutionary dynamics observed in other terrestrial small mammal reservoirs, where rodent-borne betacoronaviruses (subgenus *Embecovirus*) exhibit remarkably low barriers to cross-species transmission, frequent recombination, and an absence of host-phylogenetic structuring among sympatric species (Kotwa et al., 2026). The evolutionary constraints operating on bat-associated *Ambecovirus* therefore appear fundamentally more restrictive, reinforcing the hypothesis that chiropteran hosts maintain highly stable, species-specific relationships with their respective *Betacoronavirus* lineages (Kotwa et al., 2026).

Ultimately, the evolution of Neotropical alphacoronaviruses is best explained by a dynamic combination of long-term, stable host-virus associations punctuated by episodic cross-species transmission (CST) events, with the family Phyllostomidae serving as the central evolutionary reservoir (Caraballo, 2022; Wallau et al., 2025). On one hand, our findings strongly expand on this pattern of host-associated structuring. This is clearly evidenced by our recovery of *Amalacovirus*, *Amalacovirus*-like, and novel deeply divergent lineages actively circulating within specific phyllostomid genera, namely *Artibeus*, *Carollia*, *Glossophaga*, *Lonchorhina*, and *Phyllostomus*, reflecting a history of co-divergence where viruses accumulate genetic differences alongside their hosts over evolutionary time. Notably, our findings also reveal a clear biogeographical partitioning of these viral lineages across the studied biomes. While the novel candidate subgenus *Phyllacovirus* was recovered exclusively from the Atlantic Forest of Rio de Janeiro (FMA520 and FMA536), the *Amalacovirus*, *Amalacovirus-*like (NAC28, NAC60, PC112, and PC122), and *Phyllobecovirus* (NAC39) genomes were all detected within the semi-arid Caatinga of Ceará. This spatial distribution suggests that biome-specific environmental conditions and localized bat assembly dynamics may play a critical role in shaping the geographic distribution and maintenance of distinct coronavirus lineages in the Neotropics. On the other hand, the remarkable ecological and trophic diversity of Phyllostomidae, encompassing frugivorous, nectarivorous, insectivorous, and omnivorous species, creates high sympatric overlap in Neotropical forests (Luis et al., 2013; Jones et al., 2023). This close ecological proximity facilitates frequent contact, driving localized host-switching events almost exclusively within the family boundaries (Caraballo, 2022). Thus, the evolutionary history of these viruses in the Neotropics is characterized by relatively stable host relationships periodically interrupted by localized spillover events among sympatric bat species.

Our findings also carry critical implications for genomic surveillance and public health under a One Health framework. As anthropogenic land-use changes continue to disrupt Neotropical ecosystems, they alter bat community structures and create novel spillover interfaces. This ecological dynamic is illustrated by Jiménez-Rico *et al*. (2026), who demonstrated that lower-diversity, human-modified habitats select for generalist bat species harboring coronaviruses with wider host ranges and higher host phylogenetic diversity. In Brazil, this zoonotic risk is exemplified by the recent detection of MERS-related coronaviruses (*Betacoronavirus cameli*, subgenus *Merbecovirus*) in synanthropic *Molossus molossus* bats roosting in human dwellings in southern regions (Sita et al., 2026). The high-density gregarious behavior of synanthropic bats in periurban areas, combined with our discovery of a diverse array of novel Alpha and Betacoronavirus species circulating in wild phyllostomids across distinct biomes, highlights the urgent need for sustained, integrated genomic eco-vigilance at the human-wildlife-livestock interface.

The study presents some limitations. As this study employed an RNA-based metatranscriptomic approach, the natural degradation of RNA may have influenced the recovery of viral sequences. In addition, the samples subjected to metatranscriptomic sequencing had previously undergone freeze–thaw cycles for Sanger sequencing analyses in another study, which may have contributed to a reduction in the quality or quantity of the available RNA and, consequently, affected the recovery of viral reads. The sensitivity of the metagenomic approach may also result in the detection of viral sequences unrelated to the primary objective of the study, reflecting the natural complexity of the virome present in the samples. Finally, the costs associated with reagents and sequencing limit the possibility of repeating library preparation and sequencing experiments.

## 4. Methods

### 4.1. Sampling and RNA Sequencing

Bats were sampled from two distinct regions of Brazil. Detailed information on sampling origins has been published elsewhere (Da Silva Junior et al., 2025; Figueiroa et al., 2025). Briefly, sampling occurred from October 2020 to September 2022 in the Atlantic Forest biome within the Pedra Branca Forest (PBF), Rio de Janeiro (Da Silva Junior et al., 2025), and from March 2021 to March 2022 in the Northeast region of Brazil, Ceara State (Figueiroa et al., 2025). Bats were captured using ground-level mist nets. The mist nets were kept open for four to five hours following sunset. At the field laboratory, each captured bat was identified to the species level, and then sampled using oral and rectal swabs, and fecal samples were collected when available. All biological material was preserved in 1 mL of RNAlater™ Stabilization Solution (Invitrogen™) for subsequent laboratory analyses.

The project was authorized by the Chico Mendes Institute for Biodiversity Conservation (ICMBio) under permits 19037-1, 13373-1, 60058, and 84226; by the Environmental Institute of Rio de Janeiro State (INEA) under permit INEA020/2011; and by the National System for the Management of Genetic Heritage and Associated Traditional Knowledge (SisGen) under registrations AFAC9D5, AAB9E36, A46B0E1, A344FB0, and A03F9E2. All procedures followed the guidelines of the Ethical Committee on Animal Use of the Oswaldo Cruz Foundation (CEUA/Fiocruz; protocols LM6-18, L-36/18, LW57-19, and LW58-19) and the State University of Ceará (UECE; protocol CEUA 10802574/2021).

RNA from each sample was extracted using the QIAamp Viral RNA Mini Kit (Qiagen) according to the manufacturer’s protocol. All captured bats were screened for coronaviruses using a pan-coronavirus RT-PCR assay targeting the RNA-dependent RNA polymerase (RdRp) gene, following the protocol described by (Chu et al., 2011). The RNA from the positive sample was subsequently treated with DNase using the TURBO DNA-free™ Kit (Ambion, Thermo Fisher Scientific, Waltham, MA, USA) to remove residual genomic DNA. A total of 30 µL of RNA was mixed with 3 µL of 10× TURBO DNase buffer and 1 µL of TURBO DNase and incubated at 37 °C for 30 min. DNase activity was inactivated by adding 3 µL of inactivation reagent, followed by 5 min of incubation at room temperature. The reaction was centrifuged at 10,000 × g for 1.5 min, and the clarified supernatant was transferred to a new tube.

Host rRNA depletion and library preparation were performed using the Illumina Stranded Total RNA Prep, Ligation with Ribo-Zero Plus kit (Illumina, San Diego, CA, USA). The workflow includes enzymatic RNA fragmentation, adapter ligation, and strand-specific cDNA synthesis. Libraries were sequenced on an Illumina NextSeq platform using a NextSeq 1000/2000 P2 flow cell 100 cycles.

### 4.2. Metagenomic assembly

The viral metagenomic analysis was performed using an in house workflow. Briefly, raw paired-end FASTQ files were first processed with fastp v0.23.2 (S. Chen et al., 2018) for adapter trimming and quality filtering with the following parameters: a minimum read length of 36 bp, a phred quality score threshold of 20 (--qualified_quality_phred 20 -l 36), and automatic adapter detection (--detect_adapter_for_pe). Trimmed reads from each sample were assembled de novo using the SPAdes v3.15.0 assembler (Prjibelski et al., 2020) with --meta mode. To ensure sample traceability, contig headers were prefixed with sample-specific identifiers, and contigs from all libraries were combined for viral identification.

All assembled contigs were then subjected to viral identification by a two-stage similarity search strategy as followed by da Silva et al. (2024): first, a similarity search against a curated custom viral protein database using DIAMOND v2.0.11 (Buchfink et al., 2015) on blastx mode and --more-sensitive parameter reporting the top 100 alignments per query. The database was composed of all viral proteins from NCBI (TaxID: 10239) and deduplication was performed by clusterization with CD-HIT (Li and Godzik, 2006) to remove sequence redundancy at 95% similarity. In order to enhance the detection of novel, highly divergent, or unclassified viruses, this deduplicated dataset was supplemented with curated viral RdRp collections from NeoRdRp (Sakaguchi et al., 2022), PalmDB (Edgar et al., 2022), RVMT (Neri et al., 2022), and RdRp-Scan (Charon et al., 2022). Contigs that have matched with a viral sequence in the first similarity search were further analyzed for a confirmatory viral origin and to exclude false positives. The second DIAMOND search was performed against the complete NCBI non-redundant (NR) database, retaining exclusively those contigs consistently classified as viral in both searches. DIAMOND blastx analysis was performed with the same parameters of the first one; the five top hits were selected based on the highest bitscore, lowest e-value, and highest identity. If at least one of the 5 best hits represent a virus the contig was then treated as a viral contig. Confirmed viral contigs were analyzed with viralComplete to evaluate viral genome completeness (https://github.com/ablab/viralComplete). Taxonomic assignment was resolved using an in-house Python script that parses NCBI Taxonomy dump files (nodes.dmp and names.dmp). By tracing each retrieved TaxID back to the root of the taxonomic tree, the corresponding taxonomic hierarchy was obtained, including superkingdom, phylum, class, order, family, genus, and species.

Scaffolding of fragmented genomes were performed, recovering contigs with taxa ID assigned as *Alphacoronavirus* genus or as belonging to an unassigned genus within the *Coronaviridae* family. These sequences were then subjected to BLASTn searches (Altschul et al., 1990) against the NCBI Virus RefSeq database (default parameters, search performed on 11 November 2025). Contigs from the same sample that matched the same viral species were subsequently processed using the Multi-CSAR tool (K.-T. Chen et al., 2018), employing the best complete or draft reference genome from BLASTn analysis.

Genomes considered complete were analyzed with Pyrodigal v3.7.0 (Larralde, 2022) with default parameters to predict coding regions. Open reading frames (ORFs) predicted by Pyrodigal were submitted to a BLASTp analysis using the PSI-BLAST algorithm on the NCBI platform against viral proteins (ClusteredNR database, updated on 17 January 2026).

### 4.3. Phylogenetic Analysis

In order to reconstruct the evolutionary history of the *Alpha* and *Betacoronavirus* genomes identified with metagenomic analysis we used the same dataset described in a recent study for *Betacoronavirus* (Wallau et al., 2025). For *Alphacoronavirus* we applied the same sampling strategy described in Wallau et al. (2025). Briefly, ORF1ab proteins were recovered from NCBI Virus (NCBI, 2026) searching for “Alphacoronavirus” and applying the “Has proteins” filter for the terms “ORF1ab polyprotein, replicase polyprotein 1ab, ORF1ab, orf1ab polyprotein, polyprotein 1ab, ORF1ab protein, 1AB polyprotein, 1ab polyprotein, polyprotein ORF1ab, polyprotein orf1ab” (accessed on 16 October, 2025). The sequences were then subsampled to reduce highly similar sequences based on Genus plus Virus name, using the same in-house script available at https://github.com/dezordi/auto-ncbi/blob/master/split_by_tax.py. Sequences with unknown subgenus were kept as well. This procedure generated a final dataset of 125 *Alphacoronavirus* ORF1ab proteins. Additionally, four *Gammacoronavirus* ORF1ab proteins were included as outgroups.

Two multiple sequence alignments - for *Alpha* and *Betacoronavirus* - were performed using MAFFT v7 (Katoh et al., 2019) with default parameters for nucleotide alignment and E-INS-i strategy for multiple domains with long gaps and BLOSUM45 Matrix for divergent proteins. Alignments were inspected using Aliview v1.30 (Larsson, 2014) and only the region of 3CLpro, NiRAN, RdRP, ZBD and HEL1 domains was kept for subsequent analysis as preconized by ICTV for the *Orthocoronavirinae* subfamily (<u>Supplementary File 1</u>, amalacovirus.aa.domains.algn.fasta).

Scaffolded and unscaffolded sequences were analysed using the ORF Finder tool. Predicted regions corresponding to ORF1ab were incorporated into each reference alignment. ORFs derived from the same scaffold or contigs that aligned to distinct regions of the reference alignment were merged and considered as a single ORF.

Phylogenetic reconstructions were conducted with IQ-TREE v2 (Minh et al., 2020) following model selection by ModelFinder (Kalyaanamoorthy et al., 2017), integrated within IQ-TREE. Branch support was assessed using the approximate likelihood ratio test (aLRT) and ultrafast bootstrap (UFboot) with 1,000 replicates. The trees were annotated with ggtree (Xu et al., 2022) using both virus subgenus and host taxonomy information from the NCBI records.

#### 4.3.1. *Ambecovirus* subgenus analysis

As the *Ambecovirus* subgenus was only recently described (Wallau et al., 2025), and no subsequent studies have reported additional genomes from this subgenus, we performed a phylogenetic analysis to contextualise the genome described in this study. The ORF1ab protein sequence was aligned to a previously published alignment comprising 1,978 amino acids, provided as supplementary material in Wallau et al. 2025. Sequence alignment was carried out using MAFFT v7 with the --add and --keep-length parameters. Phylogenetic reconstruction was performed with IQ-TREE 2, applying the same model selection strategy and branch support criteria described above.

#### 4.3.2. *Amalacovirus* subgenus analysis

Only one viral sequence belonging to the *Amalacovirus* subgenus was available in the NCBI Virus database (MT663548.1). Additionally, few sequences were found in the NCBI Protein and Nucleotide databases. Considering that, to perform a phylogenetic analysis focusing on *Amalacovirus* clade, a BLASTp analysis was performed on 11 November 2025 using the ORF1ab from *Amalacovirus* reference sequence (GenBank accession MT663548.1) as query. Searches were conducted against the NCBI non-redundant protein database (nr), restricted to the *Coronaviridae* family and excluding *SARS-CoV-2*. Parameters were set to max_target_seqs = 500, word_size = 3, and substitution matrix = BLOSUM50.

For each organism returned in the search, the hit with the highest similarity to the query sequence was retained. The same procedure was applied to the ORF1ab gene from each of the six sequences previously identified within the *Amalacovirus* clade in the initial phylogenetic analysis.

All retrieved hits were concatenated, resulting in a total of 517 sequences. Duplicate entries, defined as sequences sharing both identical headers and identical amino acid sequences, were removed using seqkit rmdup -s, eliminating 288 duplicates. The final dataset comprised 229 unique sequences, which were subsequently included in the multiple sequence alignment used for the reconstruction of the *Alphacoronavirus* phylogeny.

Complementarly, a secondary phylogenetic analysis was conducted using sequences described as *Amalacovirus* for different studies. To do it, we performed a search using the “*Amalacovirus*” term on NCBI PubMed and PMC (search as performed on 6 December 2025) which returned 7 studies (details on Supplementary Table 1). Three studies only mentioned “*Amalacovirus*” in the introduction, and four studies described *Amalacovirus* clades into phylogenetic analysis. The sequences described as *Amalacovirus* in each study were recovered and used to create a curated dataset of 98 genetic fragments (446 nucleotides) of viruses that group with the MT663548.1 genome as well as that forms a sister-clade of *Amalacovirus*.

These fragments and the correspondent fragments of six *Alphacoronavirus* genomes produced in this study were used as a query into a BLASTn analysis against NCBI *Coronaviridae* family database and Zoover database (Liu et al., 2026) (update on 11 December 2025). The hits are treated to remove redundancy (by accession id since Zoover use sequences from NCBI), which returned 4,204 sequences that were clusterized with cd-hit-est using 95% of identity as threshold, forming 457 clusters that are concatenated with the curated dataset from literature and with the sequences produced in this study, generating a dataset of 560 sequences (<u>Supplementary File 1</u>, amalacovirus.nt.fragment.algn.fasta). The sequences were then aligned with MAFFT v7 with default parameters and a phylogenetic tree was reconstructed with IQTREE-2 using the same logic of model selection and branch support described above.

### 4.4. Similarity Analysis

To assess the degree of similarity between the newly generated genomes and representative viruses of related clades, we performed similarity analyses of the whole genomes and of the 3CLpro, NiRAN, RdRp, ZBD, and HEL1 domains for *Amalacovirus* and *Ambecovirus*. For each genus, we restricted the comparisons to the sequences of the most closely related clade recovered in the phylogenetic analyses for which complete genomes were available, to avoid underestimations based on non sequenced regions. Accordingly, the *Amalacovirus* dataset comprised three sequences (the two genomes generated here, FMA_520 and FMA_536, and the closest complete reference, Bat alphacoronavirus AMA_L_F, MT663548.1), whereas the *Ambecovirus* dataset comprised three sequences (NAC39, generated here, and the closely related N107 sequences from Pernambuco, Brazil (Wallau et al., 2025). Nucleotide and amino acid sequences were retrieved from the alignments described above to generate sub-alignments, which were then analysed with CIAlign v1.1.4 (Tumescheit et al., 2022) using the parameters --make_similarity_matrix_input, -- make_simmatrix_keepgaps 0, --make_simmatrix_dp 4, and --make_simmatrix_minoverlap 1.

### 4.5 Recombination

To understand whether the genomic architecture of the two novel *Alphacoronavirus* and the new genomes generated in this study (FMA_520 and FMA_536) is the result of recombination, we assembled a multiple sequence alignment comprising these genomes together with 135 publicly available *Alphacoronavirus* genomes retrieved from NCBI. Considering that recombination-detection methods are highly sensitive to alignment artefacts, the raw alignment was curated with CIAlign v1.1.4 (Tumescheit et al., 2022) using the -- remove_insertions and --crop_ends functions under default parameters (insertions of 3–200 bp flanked by at least 5 conserved positions and present in fewer than 50 % of the sequences were removed; alignment ends were cropped where the proportion of gaps exceeded 0.05, with a redefinition threshold of 0.1), followed by the removal of gap-only columns. This procedure removed poorly aligned, taxon-specific insertions and ragged terminal regions, reducing the alignment to 29,008 residues. Five low-quality genomes were further excluded manually resulting in a final alignment of 132 genomes. Finally, all ambiguous IUPAC nucleotide codes (R, Y, K, M, S, W) remaining in 39 sequences were recoded as N, since the recombination-detection algorithms implemented in OpenRDP (including GENECONV v1.81a) only accept unambiguous nucleotide characters.

Recombination analysis was performed in RDP5 (v5.93) (Martin et al., 2021). Three complementary approaches were applied. First, the Pairwise Homoplasy Index (PHI) test was used to assess the overall recombination signal across the full alignment. Second, an exploratory recombination scan was conducted in RDP5 by combining seven detection algorithms (RDP, GENECONV, BootScan, MaxChi, Chimaera, SiScan, and 3Seq). Recombination events were considered reliable when detected by at least three methods with an associated p-value below 0.05. Finally, the same detection settings were applied to a targeted analysis of FMA_520 and FMA_536 genomes, with the “Auto mask for optimal recombination detection” option enabled.

## Funding

The Brazilian National Council for Scientific and Technological Development CNPq, 303643/2022-6, 304355/2018-6, 402457/2020-0, Oswaldo Cruz Institute, Carlos Chagas Filho Foundation for Supporting Research in the State of Rio de Janeiro FAPERJ, 010.002152/2019, 260003/002694/2020, E-26/010.001597/2019, E-26/204.243/2021, E-26/200.395/2022, E-26/200.631/2022, Reference Laboratories from Oswaldo Cruz Foundation. Fundação Cearense de Apoio ao Desenvolvimento Científico e Tecnológico (FUNCAP, no 06347980/2020). G.L.W. hold fellowships from Conselho Nacional de Desenvolvimento Científico e Tecnológico (Grant processes 307209/2023-7).

## CRediT authorship contribution statement

**Leonardo Corrêa da Silva Junior:**Conceptualization, Methodology, Investigation, Writing - Original Draft, Writing - Review & Editing, Data curation. **Filipe Zimmer Dezordi:** Methodology, Software, Formal analysis, Data curation, Writing - Original Draft. **Alexandre Freitas da Silva:** Writing - Original Draft, Software, Formal analysis. **Jennifer Oliveira Melo:** Writing - Original Draft, Writing - Review & Editing. **Yago José Mariz Dias:** Writing - Original Draft, Software, Data curation, Formal analysis. **Paola Cristina Resende:** Resources, Funding acquisition. **Luciana Reis Appolinario:** Conceptualization, Methodology, Investigation, Original draft. **Marilda Siqueira:** Project administration, Funding acquisition. **Maria Ogrzewalska:** Supervision, Validation, Original draft, Project administration, Funding acquisition. **Gabriel da Luz Wallau:** Writing - Original Draft, Supervision, Methodology, Software, Formal analysis, Investigation, Data curation.

## Data availability

The genomes generated in this study are available on NCBI (Accession Numbers: QB009908-QB009913 for Alphacoronavirus and PZ946991 for Betacoronavirus). Raw sequencing reads were deposited in the NCBI Sequence Read Archive (SRA) under BioProject accession [PRJNA1455808] (accession numbers [SRR40344209–SRR40344215]).

## Acknowledgments

We thank the Platform of Instituto Oswaldo Cruz, Fiocruz, Rio de Janeiro, Brazil, and Genomic Platform DNA Sequencing RPT01A (Rede de Plataformas Tecnologicas Fiocruz). To the curator of the mammal collection at the MHNCE Aldo Caccavo de Araújo. To the owners of the locations where we carried out the fields, as well as everyone who helped during this period: Adria Teles, Artur Júnior, Beatriz Almeida, Clarissa Nobre, Claudia Velasquez, Felipe Pessoa, Hugo Fernandes, Igor Gutierrez, Karlla Morgana, Luiz Carlos, Marco Crozariol, Mateus Duarte, Roberio Freire, Rodrigo Gonzalez, Sheila Fernandes and Thabata Cavalcante.

## Notes

### Competing Interest Statement

The authors have declared no competing interest.

